# Automating Three-dimensional Osteoarthritis Histopathological Grading of Human Osteochondral Tissue using Machine Learning on Contrast-Enhanced Micro-Computed Tomography

**DOI:** 10.1101/713800

**Authors:** S.J.O. Rytky, A. Tiulpin, T. Frondelius, M.A.J. Finnilä, S.S. Karhula, J. Leino, K.P.H. Pritzker, M. Valkealahti, P. Lehenkari, A. Joukainen, H. Kröger, H.J. Nieminen, S. Saarakkala

## Abstract

**Objective:** To develop and validate a machine learning (ML) approach for automatic three-dimensional (3D) histopathological grading of osteochondral samples imaged with contrast-enhanced micro-computed tomography (CEμCT).

**Design:** Osteochondral cores from 24 total knee arthroplasty patients and 2 asymptomatic cadavers (n = 34, Ø = 2 mm; n = 45, Ø = 4 mm) were imaged using CEμCT with phosphotungstic acid-staining. Volumes-of-interest (VOI) in surface (SZ), deep (DZ) and calcified (CZ) zones were extracted depthwise and subjected to dimensionally reduced Local Binary Pattern-textural feature analysis. Regularized Ridge and Logistic regression (LR) models were trained zone-wise against the manually assessed semi-quantitative histopathological CEμCT grades (Ø = 2 mm samples). Models were validated using nested leave-one-out cross-validation and an independent test set (Ø = 4 mm samples). The performance was assessed using Spearman’s correlation, Average Precision (AP) and Area under the Receiver Operating Characteristic Curve (AUC).

**Results:** Highest performance on cross-validation was observed for SZ, both on Ridge regression (ρ = 0.68, p < 0.0001) and LR (AP = 0.89, AUC = 0.92). The test set evaluations yielded decreased Spearman’s correlations on all zones. For LR, performance was almost similar in SZ (AP = 0.89, AUC = 0.86), decreased in CZ (AP = 0.71→0.62, AUC = 0.77→0.63) and increased in DZ (AP = 0.50→0.83, AUC = 0.72→0.72).

**Conclusion:** We showed that the ML-based automatic 3D histopathological grading of osteochondral samples is feasible from CEμCT. The developed method can be directly applied by OA researchers since the grading software and all source codes are publicly available.

## Introduction

Conventional microscopic histopathological grading of osteochondral tissue is the gold standard for assessment of osteoarthritis (OA) severity *ex vivo*. The most commonly used OA grading methods are OARSI^1^ and Mankin^2^ scoring systems^3^. Mankin scoring system was developed based on late-stage OA samples, having limitations for assessment of early OA^4^ and disease extent^5^. Consequently, the OARSI grading system was introduced later to address these issues, offering more sensitivity to the mild and moderate progressive changes in articular cartilage, as well as functional information on cartilage properties^6^. Generally, histopathological grading methods sensitive to early changes are highly valuable for drug development and basic OA research^7^. Furthermore, sensitive grading methods might potentially be utilized in developing biomarkers, which are essential when developing prevention of the late-stage disease or non-surgical disease-modifying treatments^8,9^.

The conventional histopathological methods are complex, destructive and time consuming^4^, and also unable to capture all of the OA-induced changes within the full sample volume. Recently, methods combining multiple thin sections into 3D volume through image registration have been proposed^10,11^. However, such approaches can only avert partly the problem of two-dimensionality with the expense of a more laborious protocol.

Multiple 3D histopathological grading methods for different tissues have been proposed in the literature, based on magnetic resonance imaging (MRI)^12–15^, optical imaging^16^, ultrasound^17^, and atomic force microscopy^18^. 3D grading methods could possibly serve as a reference for clinical 3D modalities, as well as higher resolution 3D techniques. Contrast-enhanced micro-computed tomography (CEμCT) has shown potential in fast quantitation of osteochondral features while preserving the sample and reducing user bias^19^. We recently introduced a protocol for contrast-enhanced micro-computed tomography (CEμCT) using phosphotungstic acid (PTA) as a collagen-specific contrast agent^20,21^, and consequently, developed a 3D OA grading system to assess each articular cartilage (AC) zone separately^22^. However, the current 3D μCT grading system still requires manual assessment, thus, having a risk for user-dependent bias. The automation of this process could provide more objective evaluations.

Recently, methods for the quantitative 3D analysis of AC surface^23,24^, calcified cartilage^25^ and full cartilage tissue^19^ degeneration, as well as chondrocyte organization^26,27^ with CEμCT, have been reported. However, most of the current methods are either limited to a single osteochondral zone^23–25^ or not validated via independent testing^19^. The current implementations could be improved by developing more generalizable methods applicable to analyze multiple different osteochondral zones while utilizing more advanced validation techniques that show their feasibility on unseen data.

The development of machine learning techniques has enabled a data-driven approach in pattern recognition and decision making without the need for explicit programming. Machine learning has been applied in clinical OA research in several domains, such as the prediction of OA severity^28–31^ and progression^15,32,33^ using X-ray radiographs^28,29,31,32^ or MRI analysis^15,30,33^. However, little attention has been paid to machine learning in pre-clinical OA research^26,34,35^.

In this study, we aim to automate the recently proposed histopathological grading^22^ of CEμCT imaged osteochondral samples using Machine Learning. The feasibility of performing the automatic grading in different cartilage zones, and the robustness of the developed models to a sample acquisition protocol change, are assessed with an independent test set.

## Materials and methods

### Sample preparation

Osteochondral cores were harvested from tibial plateaus and femoral weight-bearing areas of human knee joints. Cores were extracted from 24 total knee arthroplasty (TKA) patients and 2 asymptomatic cadavers. Samples were split into two datasets based on the core diameter:

- Cross-validation set; 19 patients, *n* = 34, Ø = 2 mm, ethical approval PPSHP 78/2013, Ethical committee of Northern Ostrobothnia’s Hospital District
- Test set; 7 patients, *n* = 45, Ø = 4 mm, ethics approval PPSHP 78/2013; PSSHP 58/2013 & 134/2015, Research Ethics Committee of the Northern Savo Hospital District

For these datasets, samples that did not contain either the cartilage or bone were excluded (*n* = 11). Detailed sample and patient distributions are given in Supplementary Table 1. After the core extraction, all the samples were kept frozen at −80°C. Before the imaging, the samples were thawed and then fixed in 10% neutral-buffered formalin for 5 days. Fixation was followed by a minimum of 8h wash in 70% ethanol and minimum 48h immersion in 70% ethanol, 1% w/v PTA solution^20,21^. To prevent sample drying during μCT imaging, each sample was wrapped in Parafilm (Parafilm M, Bemis Company Inc, Neenah, WI, USA) and orthodontic wax (Orthodontic Wax, Ortomat Hepola, Turku, Finland).

### Imaging

The imaging was conducted right after the PTA immersion was completed. Samples were imaged using a desktop μCT setup (Skyscan 1272; Bruker microCT, Kontich, Belgium; Scanning parameters: 45 kV, 222 μA, 3.2 μm voxel side length, 3050 ms, 2 frames/projection, 1200 projections, additional 0.25 mm aluminum filter).

During the imaging of the test set, we used an improved version of the data acquisition protocol by checking the sample voids – areas of deep cartilage with no PTA accumulation (supplementary video 2 in *Nieminen et al.*^20^). We observed that the voids appeared due to the insufficient diffusion time, especially in samples with very thick AC layer. In the new protocol, upon detection of a void in the μCT scan, the sample was re-immersed in PTA to allow full diffusion to deep AC.

### 3D histopathological grading

We used reconstructed data to determine the semi-quantitative 3D histopathological grades for each sample, corresponding to the analyzed zones^22^. J. Leino conducted the grading according to the previously published grading system^22^. In this study we used the following grades:

- Surface continuity: Smooth and continuous = 0; Slightly discontinuous = 1; Moderately discontinuous = 2; Severely discontinuous = 3,
- Deep cartilage (zone 3, DZ) extracellular matrix (ECM) disorganization: Normal = 0; Slightly disorganized = 1; Moderately disorganized = 2; Severely disorganized = 3
- Calcified cartilage (zone 4, CZ) ECM disorganization: Normal = 0; Slightly disorganized = 1; Moderately disorganized = 2; Severely disorganized = 3

Grade distribution is presented in Table 1 and graphically in Supplementary Figure 1. Besides the multiclass grades, we also used dichotomized grades and split them into intact/mild VOI degeneration and moderate/severe VOI degeneration groups (Grades 0 and 1 were grouped against 2 and 3).

**Table 1.** Distribution of μCT grades assessed from the reconstructions (used as ground truth). The cross-validation set contained only a small number of samples from grade 3 and a reduced number of healthy samples, while almost no healthy samples were found in the test set. Otherwise, samples were distributed relatively evenly.

| Dataset | Zone | Grade 0 | Grade 1 | Grade 2 | Grade 3 |
| --- | --- | --- | --- | --- | --- |
| Cross-validation | S | 7 | 11 | 13 | 3 |
|  | D | 8 | 16 | 8 | 2 |
|  | C | 8 | 16 | 7 | 3 |
| Test | S | 2 | 19 | 9 | 14 |
|  | D | 0 | 16 | 15 | 13 |
|  | C | 0 | 24 | 11 | 9 |

*S = Surface zone, D = Deep zone, C = Calcified zone*

### Basic data pre-processing

A python *ad hoc* software was developed to preprocess the image stacks and train the regression and classification models. The workflow of this process is illustrated in Figure 1. The reconstructed samples were loaded and oriented using the following optimization algorithm. Here, the dice score was calculated against the projection of the sample onto an XY plane and a circle fitted to the projection, aiming for maximal dice score in the optimization. The center of the sample in the XY plane was detected by finding the center of mass of the image stack summed along Z-axis (Z – sample’s depth dimension). Edges of the sample were cropped using detected center and pre-defined VOI size (1300μm·1300μm·Z for Ø = 2 mm, 2600μm·2600μm·Z for Ø = 4 mm). Orientation and edge cropping processes are further illustrated in Supplementary Figure 2.

**Figure 1.**
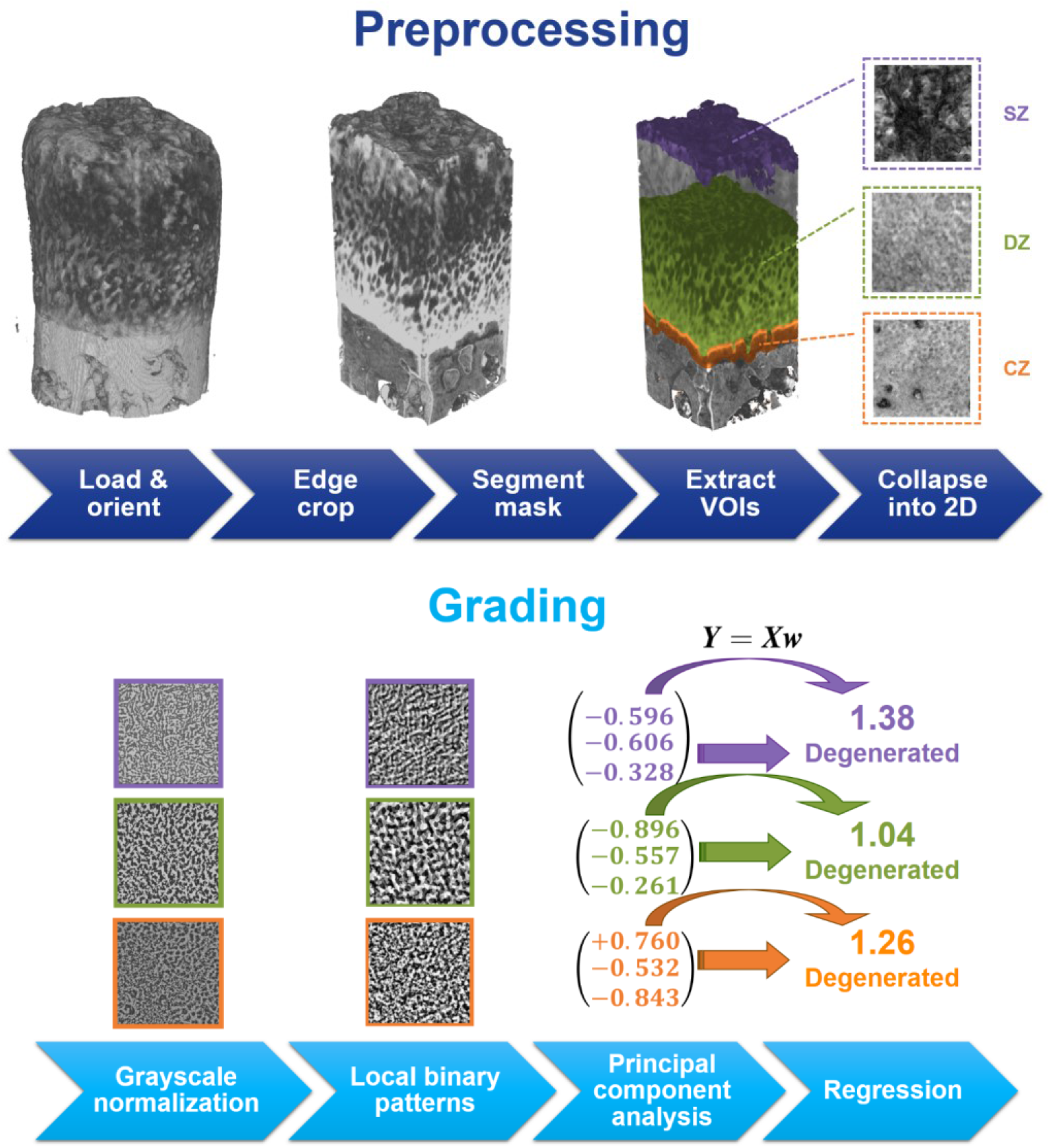
The workflow of the analysis methods used for CEμCT imaged samples. SZ = surface zone, DZ = deep zone, CZ = calcified zone.

### Calcified cartilage segmentation and VOI extraction

After cropping the sample edges, the calcified cartilage interface (tidemark) was segmented. For the Ø = 2mm samples (cross-validation set), we used the method and the pre-trained model from^36^ that allowed us to segment the calcified cartilage interface and bone automatically using U-Net – Deep Convolutional Neural Network^37^ in a slice-by-slice manner. The U-Net approach was used to consider the existing voids in the Cross-validation set during segmentation.

For the Ø = 4mm samples (test set), we used a different approach since the trained CNN model did not generalize well to a different acquisition protocol that was used for the test set data acquisition (see Supplementary Figure 3). However, the reconstructed images in the test set did not include voids and there was always a strong gradient visible at the tidemark (Figure 2a). We performed a segmentation using k-means clustering with 3 clusters. Cluster with the highest grayscale centroid belonged to the deep cartilage due to the high PTA accumulation. The area below this cluster was labeled as the calcified zone. This segmentation was performed in a slice-wise manner on XZ and YZ planes.

**Figure 2.**
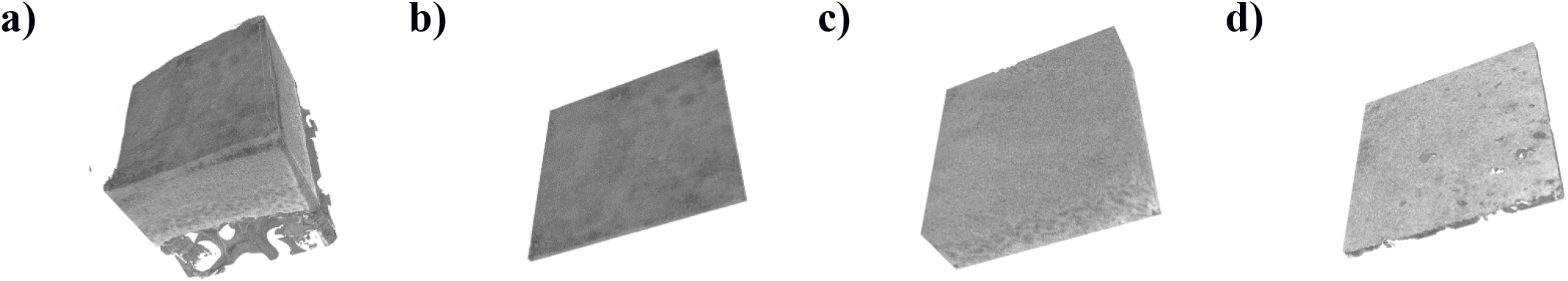
Oriented and edge-cropped VOI from a healthy / mildly degenerated osteochondral sample in the test set (harvested from an asymptomatic cadaver), **b)** Sub-VOI from the cartilage surface, **c)** deep cartilage, and **d)** calcified tissue. A smooth and continuous surface is visible. Deep and calcified ECM are well organized.

Once the calcified tissue mask was acquired, the average depth of AC was calculated using the mask and the surface coordinates of the samples. The depth for DZ was set as 60% of AC depth to ensure that the full zone was included also on delaminated samples. The lower limit for DZ was set to 30μm above the segmentation mask to ensure that the interface and calcified tissues were not included in DZ. The surface was detected using the Otsu threshold, and surface zone (SZ) was set extending 160μm below (50 slices). CZ was set as 160μm thick volume immediately below DZ. Here, we used small zone thickness values to focus on the detailed surface features and account for samples with thin CZ. Extracted volumes (Figures 2 and 3) were collapsed into two-dimensional (2D) texture images summing their mean and the standard deviation depth-wise.

**Figure 3.**
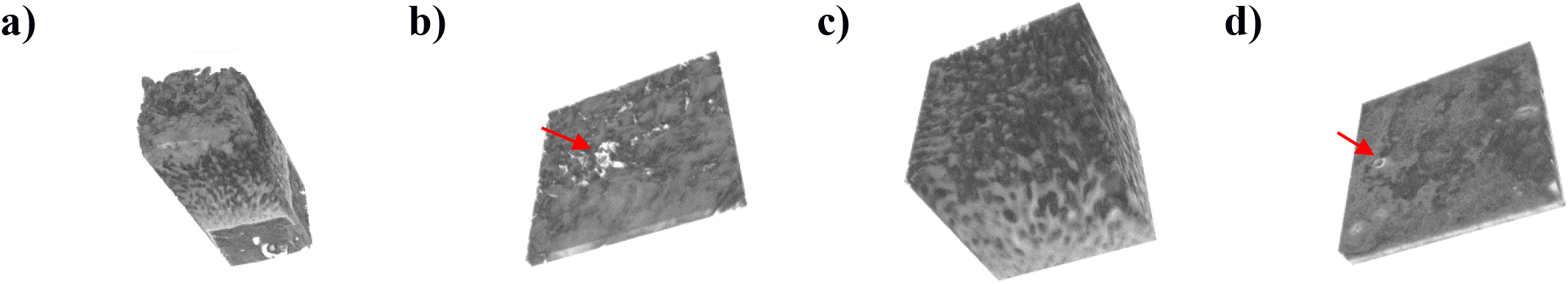
Oriented and edge-cropped VOI from a degenerated osteochondral sample in the crossvalidation set (harvested from a TKA patient), **b)** Sub-VOI from the cartilage surface, **c)** deep cartilage, and **d)** calcified tissue. Surface discontinuities, as well as deep and calcified ECM disorganization, are clearly visible. Vascular infiltration and surface discontinuities are shown with a red arrow.

Finally, all the Ø = 4mm samples included in the test set were split into nine smaller subimages (with dimensions half to the original image) to increase prediction reliability. This was also done to make sure that the textural features of the large image have similar relative size and impact on the resulting feature descriptor used to predict the 3D grades of the sample, compared to the features trained on cross-validation.

### Feature extraction

Prior to the feature extraction, possible misalignment artifacts appeared during preprocessing were automatically cropped out. In the algorithm, possible defects on the image corners were detected using adaptive thresholding and cropped. Subsequently, we performed a local normalization by subtracting from each pixel of its neighborhood’s weighted intensity. Here, we used a gaussian kernel for intensity weighing. The kernel parameters were optimized independently for each sample zone (Supplementary Table 2).

To extract the features related to cartilage degeneration, Median Robust Extended Local Binary Patterns (MRELBP) were calculated according to Liu et al^38^. Thirty-two features were extracted using rotation-invariant uniform mapping (2 from the center image, 10 from small, large and radial LBP images each). This histogram was eventually normalized to have the total sum of 1 (division by a sum of all elements). Features that did not have any occurrences were excluded resulting in 28 features. Subsequently, we mean-centered the data.

After the data centering, a principal component analysis (PCA) based whitening was used, and consequently, the dimensionality of the extracted feature vectors was also reduced. Here, 90% of the explained variance was set as a threshold for finding the optimal number of principal components. Eventually, three components were automatically selected for all the cartilage zones.

### Automatic grading

After the PCA, we used the obtained features to train two regression models on cross-validation. In particular, we used leave-one-patient-out (LOPO) cross-validation, using samples from each individual patient as a validation set, against a model trained on the rest of the patients in the dataset. The cross-validation set had two samples per patient (Supplementary Table 1). Firstly, a Ridge regression model was trained against the ground truth μCT grades. Here, we used L2 regularization with a coefficient of 0.1. Secondly, a Logistic regression model (also with L2 regularization) was trained to assess the sample’s degeneration in a binary manner.

For the test set images, the developed models were evaluated for all the nine sub-stacks separately and the average of their predictions was finally used. The models trained with the best hyperparameters from the cross-validation set were selected. To also estimate the validity of our texture-based 2D approach on the test set, separate models were subsequently trained using LOPO cross-validation (Replication experiment, see the results).

### Parameter optimization

To tune the hyperparameters for MRELBP and grayscale normalization, we used the Bayesian hyperparameter optimization algorithm from Hyperopt package^39,40^. To avoid overfitting, we performed a “nested leave-one-out” cross-validation (Figure 4). In particular, during the leave-one-out, we used a hyperparameter search on the N-1 (33 out of 34) samples using another, nested leave-one-out cross-validation. A regression model was trained for each optimization batch of 33 samples. Optimization was conducted on the cross-validation set evaluating a maximum of 100 parameter sets per iteration. The algorithm converged to the same solution on most of the iterations (30/34 for SZ, 34/34 for DZ and 18/34 for CZ) and we used the most frequent solution as the hyperparameter selection for each zone. Optimized sets of parameters are listed in supplementary table 1.

**Figure 4.**
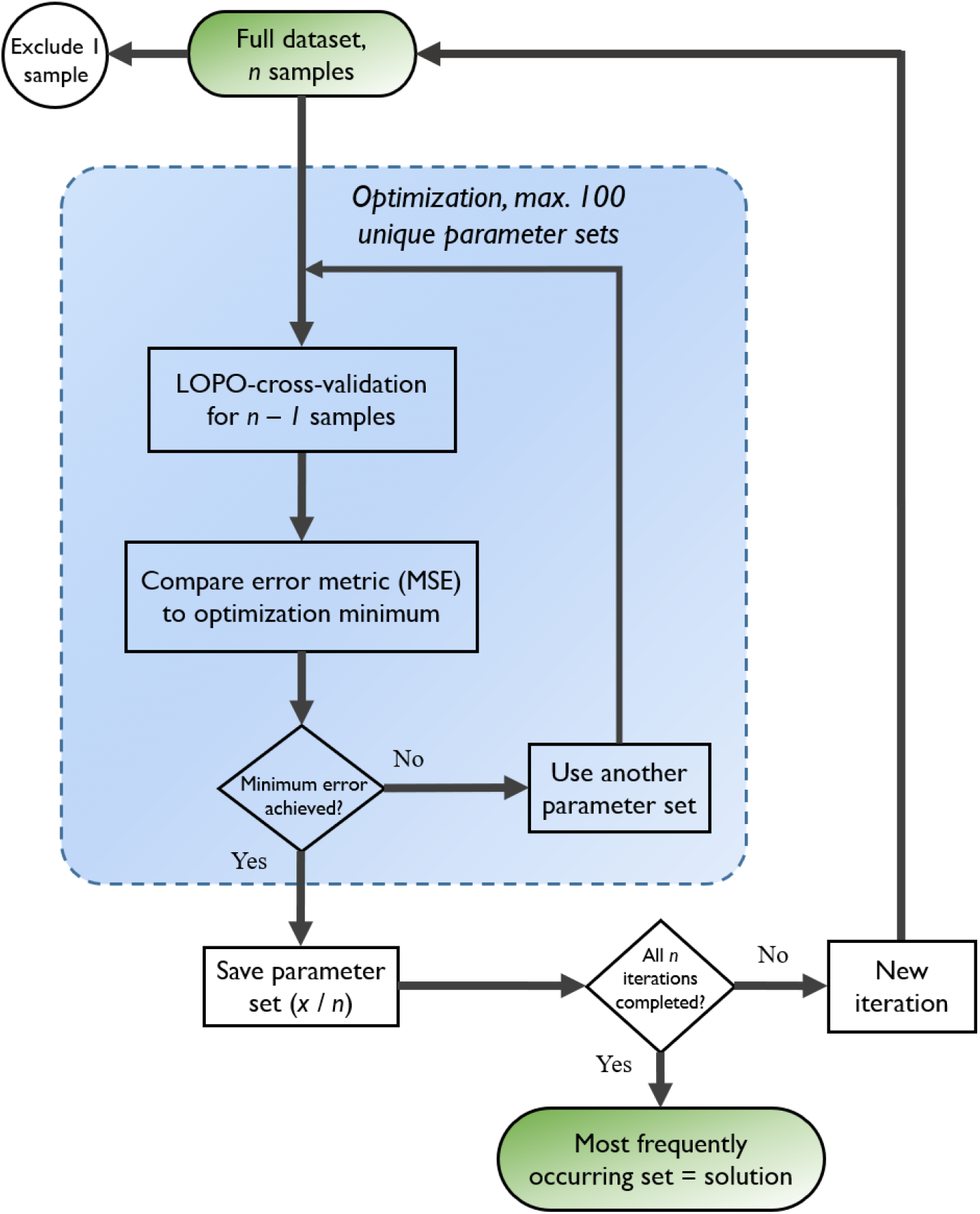
Flowchart describing the nested cross-validation method used in the parameter optimization. First, LOO is performed resulting in *n – 1* samples in the optimizations. A maximum of 100 parameter sets are evaluated in the optimization algorithm, where regression is performed with the LOPO split. Initial LOO results in 34 optimization results and the most frequent parameter set is used as a final solution.

### Statistical analyses

Predictions of the Ridge regression models were assessed using the mean squared error (MSE) and Spearman’s correlation analysis. For the Logistic regression models, receiver operating characteristic (ROC) curves and precision-recall curves (PRC) were calculated. We evaluated the area under the ROC curve (AUC) and the average precision (AP) of PRC. The 95% confidence intervals were estimated via stratified bootstrapping with 2000 iterations. To further analyze the performance of the binary classification models, we calculated the precision, recall and F1 scores under the threshold of 0.5.

## Results

### Detection of sample degeneration

For the cross-validation set, we obtained the AUCs of 0.92 (0.80, 0.99), 0.72 (0.54, 0.88) 0.77 (0.54, 0.94) for SZ, DZ and CZ, respectively. Having the threshold of 0.5 for LR’s predictions, the precision (positive predictive value) of the model was found to be high on SZ (0.83), while it remained moderate on DZ and CZ (0.44 and 0.41, respectively). The recall was found to be very high on SZ and DZ (0.94 and 0.80, respectively) and strong (0.70) for CZ. F1 scores were 0.88, 0.57 and 0.52 for SZ, DZ and CZ respectively. APs from PRC curves were 0.89 (0.77, 0.99), 0.50 (0.35, 0.75) and 0.71 (0.48, 0.91) for SZ, DZ and CZ, respectively.

For the test set, we obtained the AUCs of 0.86 (0.73, 0.95), 0.72 (0.56, 0.86) and 0.63 (0.45, 0.78) for SZ, DZ and CZ, respectively. Precisions were 0.78, 0.84 and 0.62 for SZ, DZ, and CZ, respectively. The recall was 0.61 on SZ, 0.57 for DZ and 0.40 for CZ. F1 scores for both SZ and DZ were 0.68 and 0.68, and for CZ of 0.49, respectively. APs from PRC curves were 0.89 (0.78, 0.96), 0.83 (0.73, 0.93) and 0.62 (0.48, 0.77) for SZ, DZ and CZ, respectively. Comparable detection accuracy was found for SZ compared to the cross-validation set, while a minor performance decrease was seen on CZ. The average precision of the DZ model increased by 0.33 compared to the crossvalidation set.

ROC and PRC curves (Figure 5) show that the model for SZ is performing best compared to all zones. On the cross-validation set, ROC curves show that CZ performs slightly better compared to DZ, but the difference is much more obvious in the PRC plot. Similar results can be seen on the test set, except that DZ performs better compared to CZ.

**Figure 5.**
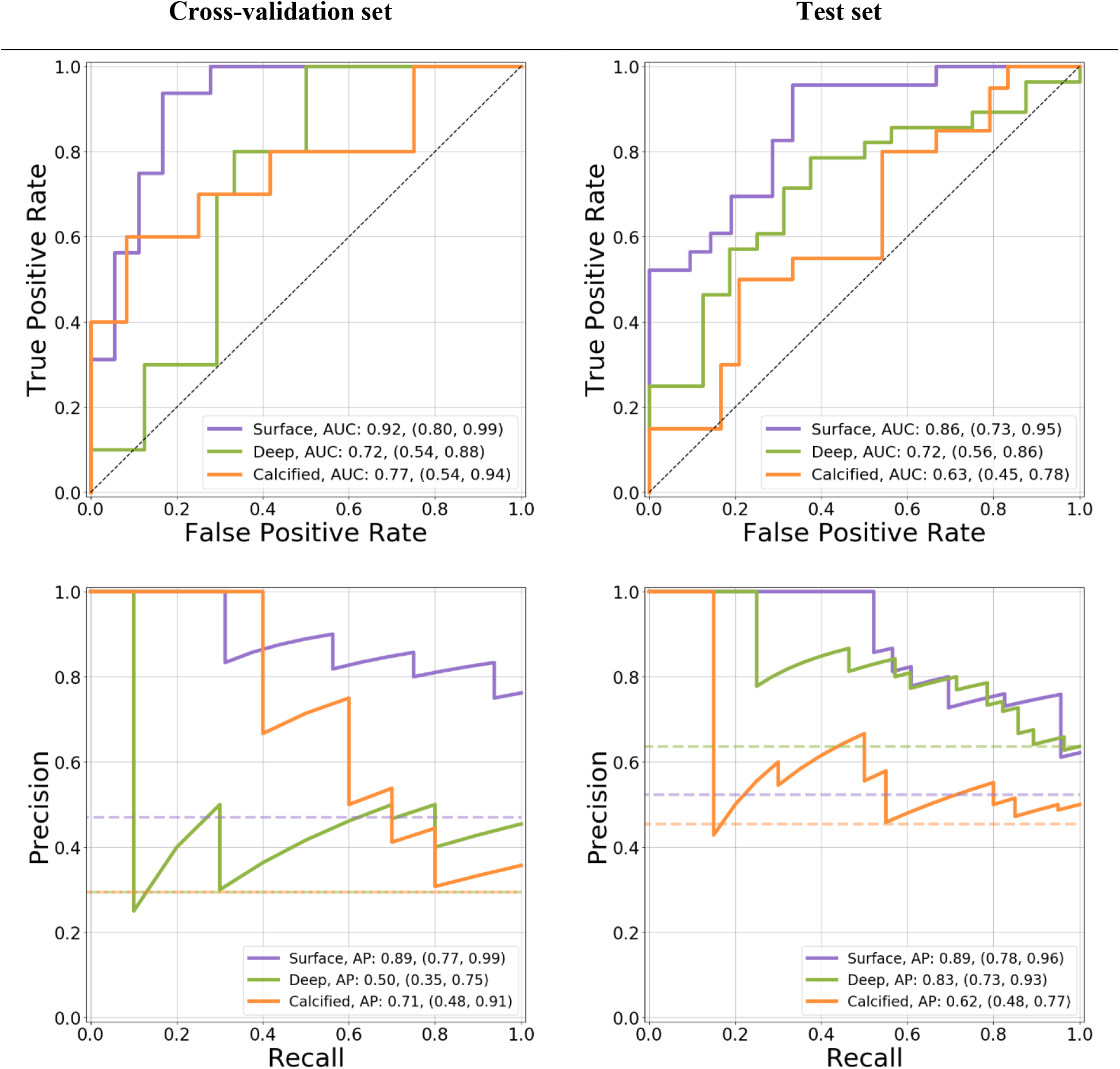
Receiver operating characteristic (ROC) and precision-recall curves (PRC) for each dataset. Values for bootstrapped AUCs and APs with 95% confidence intervals are shown. From both curves, it can be clearly seen that surface models are performing well compared to the baseline.

### Automatic grading

The performances of all the developed models are summarized in Table 2 and Figures 5–6. In particular, the Ridge regression model yielded MSEs of 0.49, 0.66 and 0.50 for SZ, DZ and CZ, respectively. Strong Spearman’s correlation was observed for SZ (ρ = 0.68), while moderate and weak correlations were observed for CZ (ρ = 0.54) on DZ (ρ = 0.38) compared to the manual grades.

**Figure 6.**
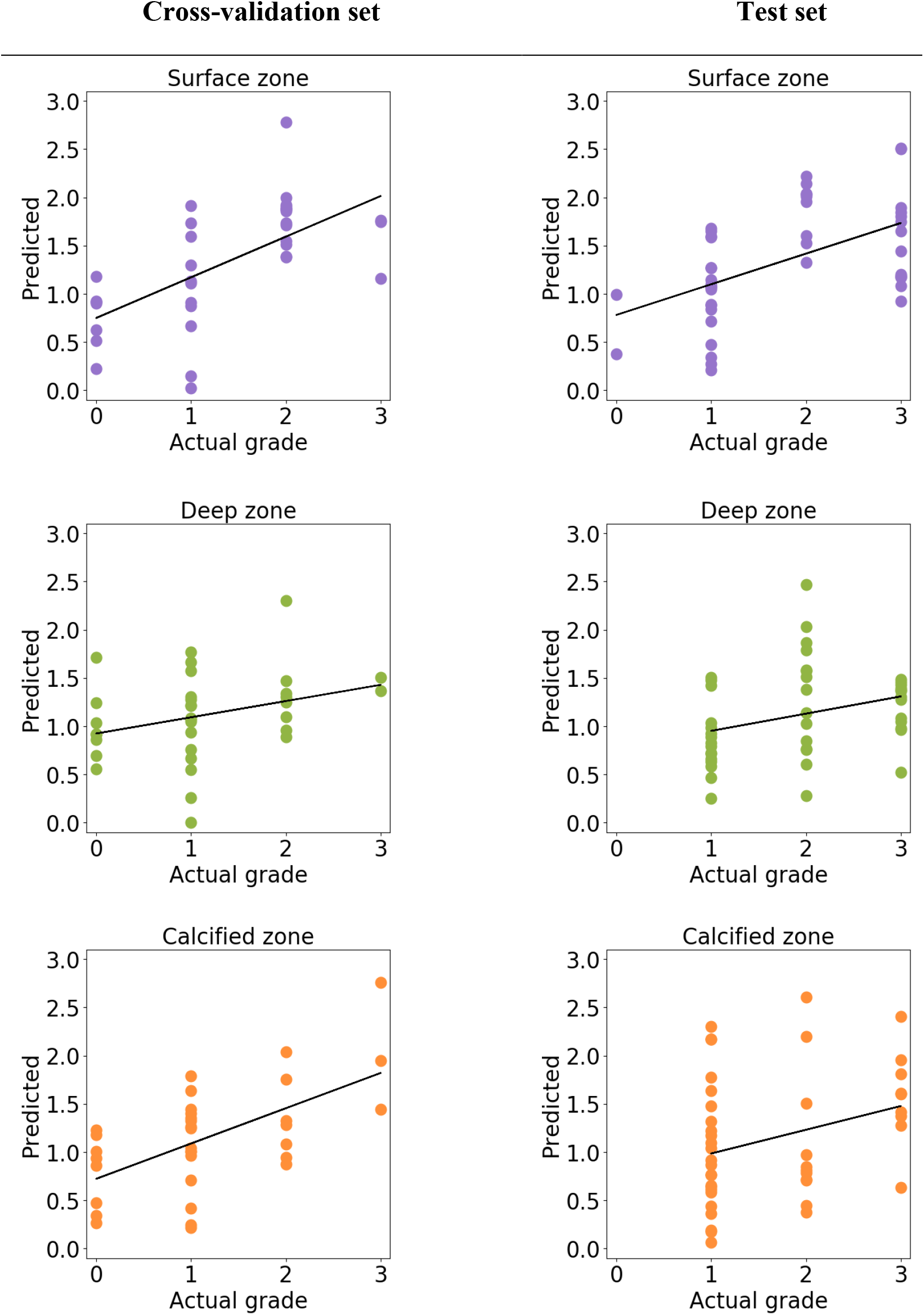
Predictions obtained from the Ridge regression models on the cross-validation (left column) and test sets (right column). Predictions in most models are very close to grade 1, showing that ridge regression has little power to distinguish individual grades in this case. On the cross-validation set, predictions for SZ and CZ as well as for test set SZ, low and high grades can be visually separated from each other.

**Table 2.**
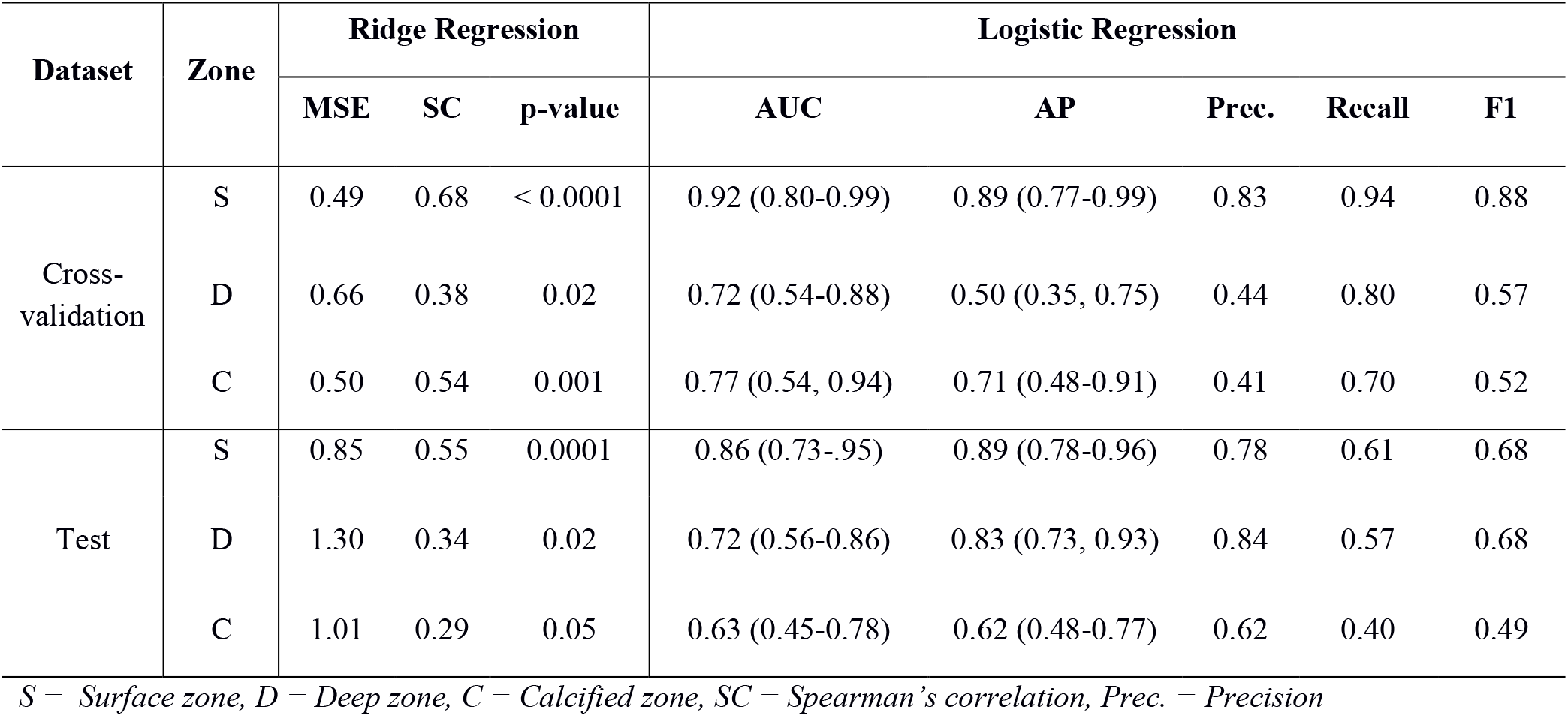
Performance of trained ridge and logistic regression models. Confidence intervals for 95% are given in parentheses. Statistical variables for ridge regression are on the left side of the table and variables for logistic regression are on the right side.

For the test set, we evaluated the predictions using the models that were saved during the training of the cross-validation set. The test set yielded MSEs of 0.85, 1.30 and 1.01 for SZ, DZ and CZ, respectively. Spearman’s correlation was moderate (ρ = 0.55) on SZ and weak (ρ = 0.34, 0.29) on DZ and CZ.

### Replication experiment

The replication experiment was performed to assess the transferability of the developed texture-based volume analysis technique. The results from the model trained separately for the test set with LOPO cross-validation are shown in Supplementary Table 3. Ridge regression showed improvement in MSE (0.85→0.69, 1.30→0.71, 1.01→0.72, for SZ, DZ and CZ, respectively) but not in Spearman’s correlation. Logistic regression yielded similar results using ROC/AUC and PRC analysis, apart from the slight increases in AUC for SZ and CZ models (0.86→0.87 and 0.63→0.64, for SZ and CZ, respectively). However, additional parameters show that recall and F1 score are improved in SZ and CZ, when using the threshold of 0.5 for the LR model (recall: 0.61→0.78 and 0.40→0.65, F1 score: 0.68→0.78 and 0.49→0.61 for SZ and CZ, respectively).

### Software Prototype

We implemented the developed automatic 3D grading method in an open-source software package for Windows OS (Supplementary Video). Currently, the models trained using a python script are exported into an intermediate format and loaded by the software to predict the degeneration of unseen samples. Additional features of the software are manual tools for artefact cropping and also the advanced visualization pipeline. The source code of the software is available on GitHub: https://github.com/MIPT-Oulu/3DHistoGrading.

## Discussion

In this paper, we investigated the feasibility of automation of the 3D μCT grading system for osteochondral human samples. We developed a method based on machine learning to predict the grades of degeneration for AC surface, deep and calcified cartilage zones in an automatic manner. The trained models were evaluated in two settings – via cross-validation and on a completely independent dataset. This allowed the assessment of generalization of the developed method to the unseen data, as well as its robustness and applicability to the new data acquisition settings.

From the experiments, we found that our models are more suited for the detection of the presence of overall degeneration in the analyzed VOI, instead of fine-grained grading. This is probably due to a limited number of training samples. However, on the other hand, this result is highly generalizable to different data acquisition settings as shown in our experiments. The results showed that the surface degeneration can be detected reliably (AUC of 0.92, F1 of 0.88 and AP of 0.89) and with moderate performance for both DZ and CZ (AUC > 0.70, F1 > 0.5 and AP > 0.5). To further increase the reliability of the presented models, novel data augmentation and semi-supervised grading techniques, *e.g.* domain adaptation^41,42^, could be utilized in the future.

On the cross-validation set, our pipeline performed better on CZ compared to DZ. However, on the test set, an AP increase of 0.33 was observed for the DZ model and a drop of 0.09 for the CZ model, respectively. Besides, during parameter optimization, the CZ model had multiple occurrences of a second parameter set. These findings suggest a better overall quality of the predictions for the DZ compared to the CZ model. The absence of the fully intact samples in the test set might be one reason for the decreased recall values when only the possibly more difficult grade 1 samples are left to be classified as negatives (Table 1, Supplementary Figure 1).

To facilitate the generalization of our method, we performed several preprocessing steps: sample preparation artefact cropping, MRELBP histogram normalization, PCA-based dimensionality reduction and whitening as well as the splitting of the larger, Ø = 4mm samples to the sub-volumes. To ensure a robust validation scheme, we used nested LOO where a Bayesian hyperparameter search was performed at each iteration of cross-validation. According to this strategy, we mitigated the risk of overfitting^43^ that is highly probable with small sample sizes.

Besides the robust validation scheme, we also tackled the issue of a thorough evaluation of the results. When making binary classification, ROC curves are often reported^44^. They are easily understood and allow assess performance well on evenly distributed datasets. However, the PRCs are more descriptive on imbalanced datasets and provide information on the positive predictive value of the models^45,46^. The use of the ROC curve analysis can even lead to false conclusions on classifier reliability when using imbalanced data due to wrong interpretations of the true positive rate^45^. We consider the use of a different metric for classification models to be one of the core strengths of this study.

Our group has previously utilized a novel method for quantitative surface morphology assessment. Similarly to the handcrafted surface features presented by *Ylitalo et al.*^23^, our machine learning approach here showed the highest sensitivity for SZ for detecting intact samples. This highlights the importance of surface features, although the presented machine learning method can provide a comprehensive description of pathological changes of other cartilage zones as well. These studies are not otherwise directly comparable either since a different split (grades 0-1 against 2-3, instead of 0 against ≥ 1) was used here to better balance the grade distributions of the different groups (class distribution in *Ylitalo et al.*^23^ was 7 against 29 for the surface). Further, in the current study, we conducted a more thorough validation with nested LOO, PRC analysis, and independent testing.

Differences in performance between the replication experiment and the experiment on the cross-validation set could be explained by the differences in the data acquisition since μCT imaging parameters were optimized for Ø = 2mm. We analyzed this both visually and quantitatively, comparing the images with the filtered data (Supplementary figure 3 and 4). For the test set, MSE against the filtered data was higher (mean MSE = 29.6) compared to the cross-validation set (mean MSE = 5.8). Both PSNR and SSIM were higher in the cross-validation set (mean values 40.2 and 0.84 compared to 33.3 and 0.71). All three metrics suggest higher data quality in the cross-validation set.

This study has several important limitations. First and foremost, a very reliable and accurate model might require hundreds or thousands of samples from different patients, and the current model was created based only on 34 samples from TKA patients. Secondly, we had to include one freeze-thaw cycle for the samples due to practical reasons. Thirdly, datasets used in the study were very heterogeneous due to different core diameters, causing decreased image quality in the test set. Fourthly, distribution of μCT grades was also different on the test set, which could be due to lower patient count or the lack of multiple graders. Finally, there are possible zone-specific limitations that should be noted: causes of error in the CZ model could be due to the use of a thin VOI or inefficient tidemark characterization by k-means clustering –based segmentation (our trained U-Net segmentation was not used for the test set since it did not generalize). Moreover, DZ model performance might increase if a smaller depth of cartilage was used (e.g. 30-40% instead of 60% of cartilage depth^47^), better avoiding inclusion of the transitional zone.

As a conclusion, this study shows that automatic 3D histopathological grading of osteochondral samples is feasible from CEμCT with minimal user input. Our model could be directly used to provide a second opinion for OA researchers requiring a reliable assessment of OA *ex-vivo* severity, especially at the surface zone. Further development of the model, including the acquisition of a bigger training dataset, would likely increase the reliability of the analysis for zones other than the cartilage surface. To the best of our knowledge, this is the first report presenting a machine learning based 3D histopathologic grading model, which also adequately generalizes to unseen data. All codes used, and the software prototype developed during this study are available on the project’s GitHub page (https://github.com/MIPT-Oulu/3DHistoGrading).

## Acknowledgements

The financial support of the Academy of Finland (grants no. 268378, and 303786); Sigrid Juselius Foundation; European Research Council under the European Union’s Seventh Framework Programme (FP/2007-2013)/ERC Grant Agreement no. 336267; and the strategic funding of University of Oulu are acknowledged.

## Author contributions

Conception and design: SJOR, AT, MAJF, SSK, HJN, SS.

Data analysis, development of the pipeline and the software prototype: SJOR, TF, AT.

Data acquisition: SJOR, MAJF, SSK, JL, MV, PL, AJ, HK.

Drafting the manuscript: SJOR, AT.

Critical revision for important intellectual content and approval of the manuscript: all authors.

## Supplementary material

**Supplementary Figure 1.**
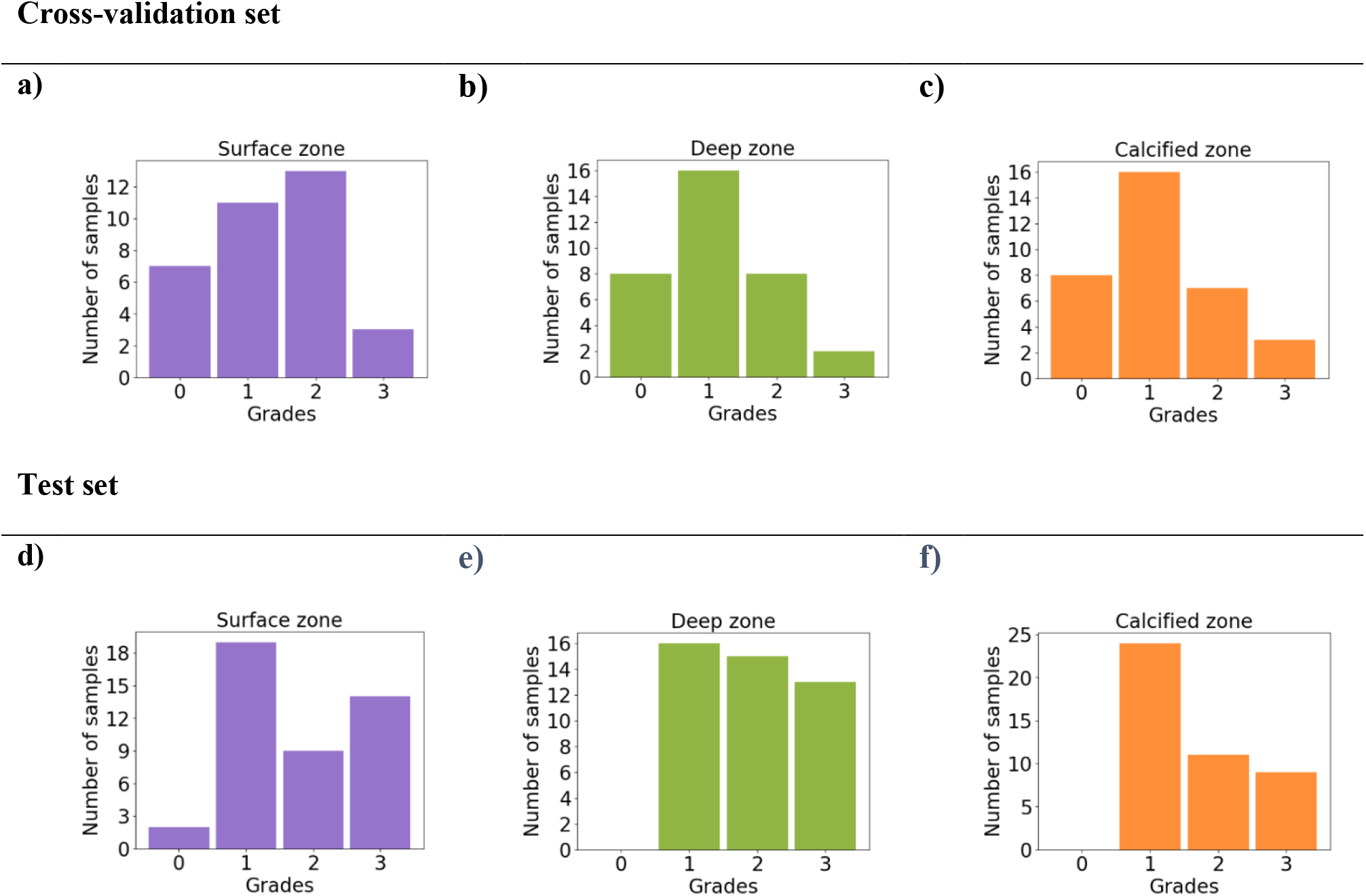
Visual representation of the grade distribution of different datasets for the tested zones. Cross-validation (**a-c**) set has the broadest distribution of μCT grades and is well suited for training the regression models (however, only small amount of grade 3’s are included). Test set (**d-f**) has almost no grade 0 samples. Exact values for the classes are listed in Table 1.

**Supplementary Figure 2.**
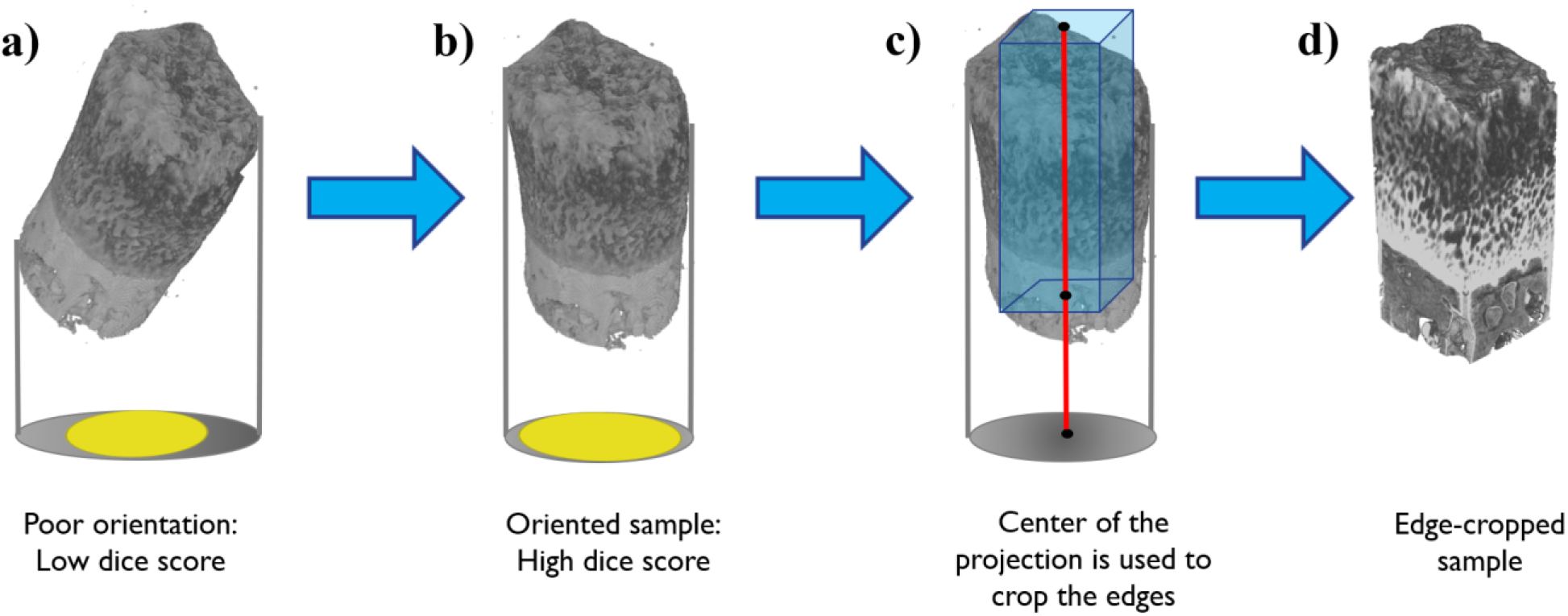
Illustration of the orientation and cropping of the samples on the preprocessing pipeline. Poorly oriented sample casts an elliptic projection along the z-axis (a). This results in low dice score against the fitted circle (yellow). Small rotations are performed in order to increase the dice score between the fitted circle and the projection (b). From the oriented sample, the center of the projection is calculated and used to crop the edges of the sample (center axis displayed in red), resulting in a rectangular cuboid VOI (blue) inside the sample (c-d).

**Supplementary Figure 3.**
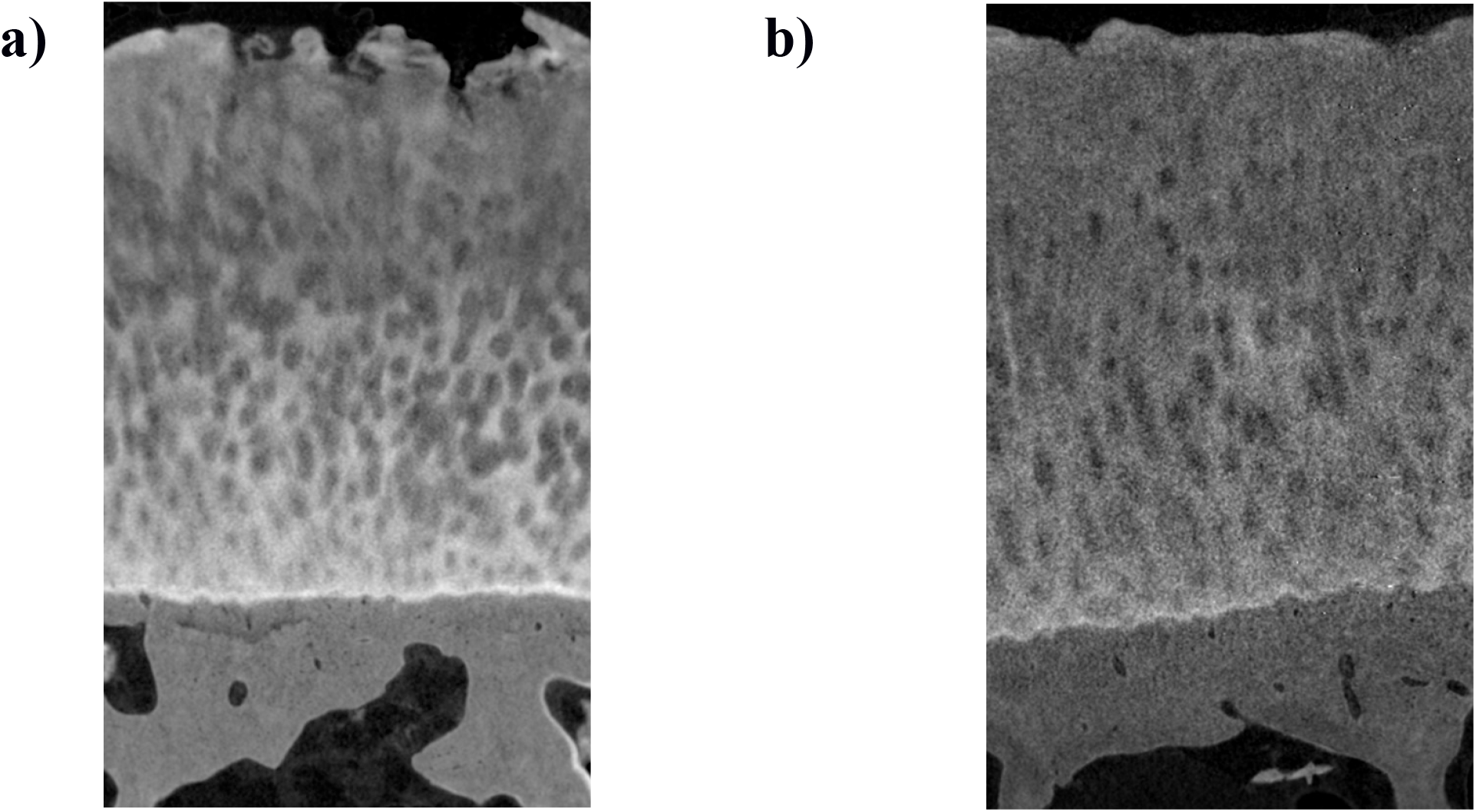
Comparison of μCT data on different datasets. In the cross-validation set (a), the core size is small, and the acquired signal is higher compared to test sets. The test set (b) has a larger core diameter, which seems to result in lower image quality due to imaging parameters optimized for the small diameter. This results in a lower measured signal on the detector. Visual differences are quantified in Supplementary Figure 4.

**Supplementary Figure 4.**
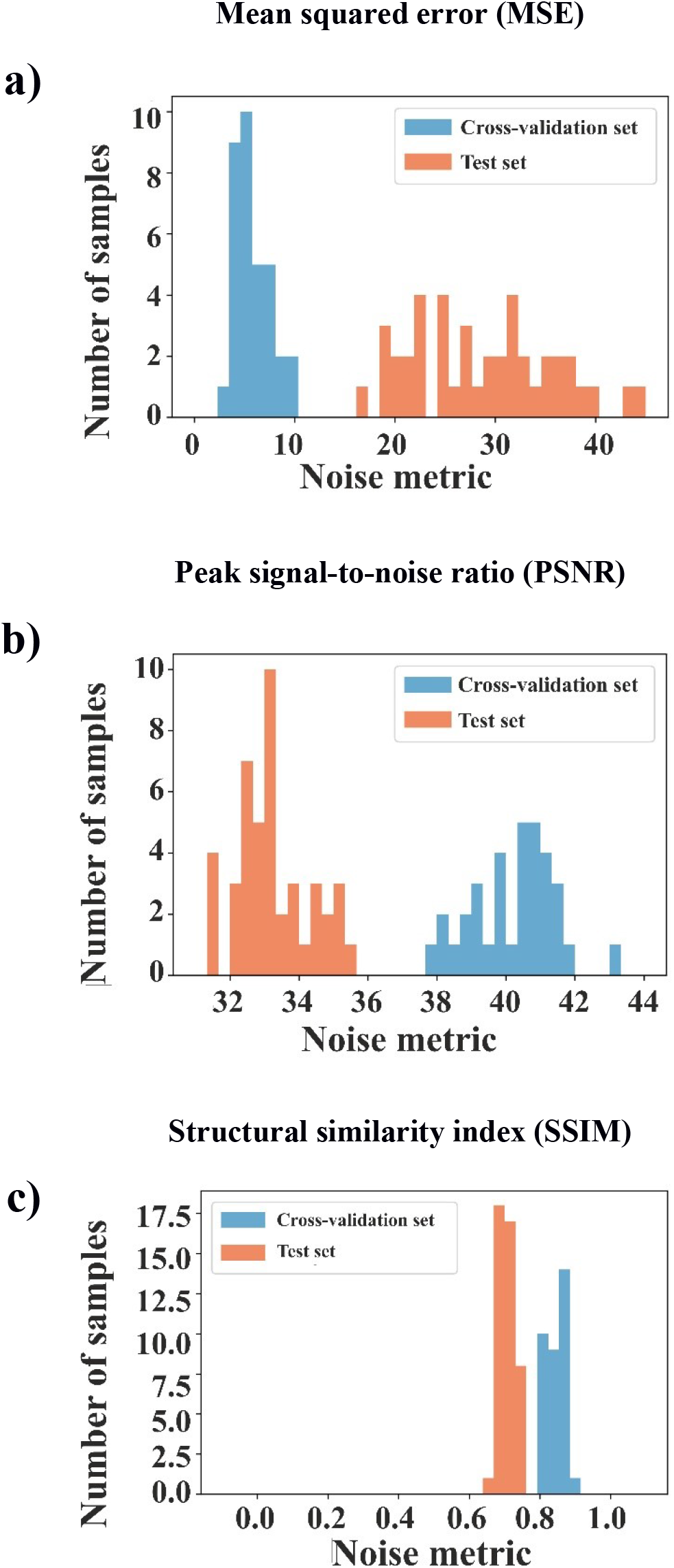
To quantify the differences in image quality of the two datasets, we calculated MSE (**a**), peak signal-to-noise ratio (**b**, PSNR) and structural similarity index (**c**, SSIM). Reconstructed coronal slices were compared against the same slices with median filtering (kernel size 5). Multiple slices were assessed along each sample to get averaged values for metrics. Histograms from individual samples are shown. Mean values for Cross-validation set are: MSE = 5.8, PSNR = 40.2, SSIM = 0.84. Mean values for test set: MSE = 29.6, PSNR = 33.3, SSIM = 0.71.

**Supplementary Table 1.**
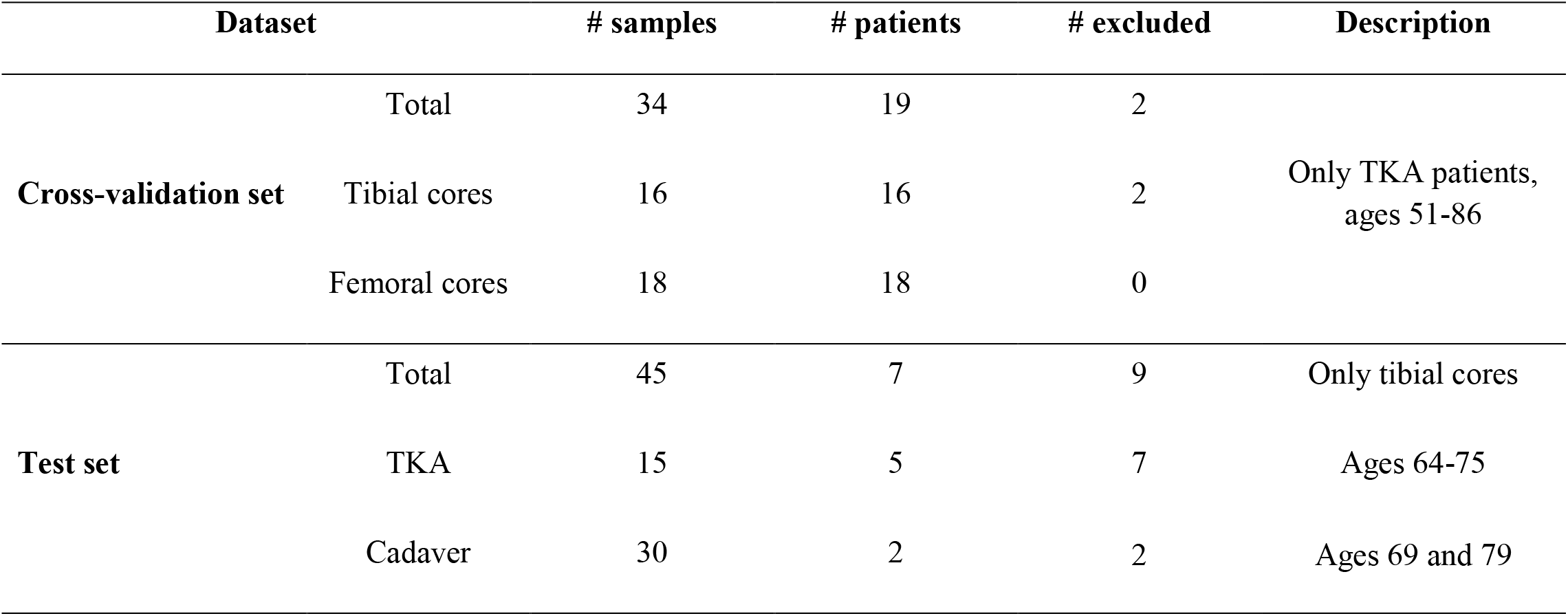
Sample and patient distribution. The cross-validation set consists of only TKA patients with one femoral and tibial sample each (four patients had a sample from only one location included). The test set consists of both cadaver and TKA patients from the tibial compartment. The number of patients is much higher on the Cross-validation set allowing large variation in training the models. Samples that were initially excluded when creating these datasets are shown (not containing either the cartilage or bone layer).

| Dataset |  | # samples | # patients | # excluded | Description |
| --- | --- | --- | --- | --- | --- |
| Cross-validation set | Total | 34 | 19 | 2 | Only TKA patients, ages 51-86 |
|  | Tibial cores | 16 | 16 | 2 |  |
|  | Femoral cores | 18 | 18 | 0 |  |
| Test set | Total | 45 | 7 | 9 | Only tibial cores |
|  | TKA | 15 | 5 | 7 | Ages 64-75 |
|  | Cadaver | 30 | 2 | 2 | Ages 69 and 79 |

**Supplementary Table 2.** Parameters optimized in contrast normalization and MRELBP with a description of each parameter.

| Parameter | Values used in all zones | Frequently encountered values in CZ (16/34) | Description |
| --- | --- | --- | --- |
| Gaussian kernel 1 | 25 | 23 | Size of the kernel for centering the input image (subtracted from input) |
| Gaussian kernel 2 | 21 | 21 | Size of the kernel for standardizing the input image (divided from image) |
| Sigma 1 | 4 | 4 | Standard deviation of Gaussian kernel 1 |
| Sigma 2 | 7 | 6 | Standard deviation of Gaussian kernel 2 |
| Neighbors | 8 | 8 | Number of neighbors used in MRELBP (4 orthogonal and 4 diagonal neighbors). |
| Large radius | 18 | 12 | Distance of center pixel from neighbors used in obtaining large image |
| Small radius | 4 | 11 | Distance of center pixel from neighbors used in obtaining small image |
| Center kernel | 15 | 9 | Kernel size used in median filtering center image |
| Large radius kernel | 15 | 9 | Kernel size used in median filtering large LBP image |
| Small radius kernel | 13 | 15 | Kernel size used in median filtering small LBP image |

**Supplementary Table 3.** Results of trained models on the test set (replication experiment). Separate models were trained using leave-one-patient-out (LOPO) cross-validation, averaging predictions from the nine substacks. Values improved due to separate training are bolded. Ridge regression shows improvement in MSE but not in Spearman correlation. The values of AUC show only slight differences in logistic regression, but additional analysis shows that recall and F1 score are improved in SZ and CZ.

| Zone | Ridge regression |  |  | Logistic Regression |  |  |  |  |
| --- | --- | --- | --- | --- | --- | --- | --- | --- |
|  | MSE | SC | p-value | AUC | AP | Prec. | Recall | F1 |
| S | **0.69** | 0.45 | <0.01 | **0.87** (0.74, 0.96) | 0.87 (0.75, 0.96) | 0.78 | **0.78** | **0.78** |
| D | **0.71** | -0.06 | 0.71 | 0.64 (0.46, 0.79) | 0.80 (0.69, 0.90) | 0.80 | 0.57 | 0.67 |
| C | **0.72** | -0.16 | 0.30 | **0.64** (0.45, 0.79) | 0.56 (0.44, 0.77) | 0.56 | **0.65** | **0.61** |

*S = Surface zone, D = Deep zone, C = Calcified zone, SC = Spearman's correlation, Prec. = Precision, F1 = F1 score*

**Supplementary Video.** Example usage of the grading and visualization software.

